# Probabilistic network modeling identifies RBPJ as a driver of stemness in Merkel cell carcinoma

**DOI:** 10.64898/2026.06.26.734761

**Authors:** Waldo Guzman Barrientos, Mireya Herrera-Herrera, Hannah Majors, Rodrigo De La Vega Gomar, Baris Kerimoglu, Megha Padi

**Author notes:** Corresponding author: Megha Padi, 1007 E. Lowell St., Tucson, AZ 85719.

## Abstract

Merkel cell carcinoma (MCC) is a poorly differentiated neuroendocrine carcinoma with limited treatment options, primarily immunotherapy, to which only ∼50% of patients respond. Lineage plasticity drives its poorly differentiated phenotype, in turn promoting tumor aggressiveness and treatment resistance. Targeting the mechanisms underlying lineage plasticity could help induce differentiation, reduce proliferation, and potentially sensitize the tumor to existing therapies, yet such strategies are underdeveloped in MCC. Here, we integrate scRNA-seq and bulk ATAC-seq data to generate Boolean networks, simulate their dynamics, and predict a key regulator of differentiation, which we validated *in vitro*. Using CytoTRACE2 across two independent datasets, we revealed the existence of tumor subpopulations with distinct developmental potency states. We then constructed and refined transcription factor regulatory networks using BooleaBayes and expanded them with ATAC-seq inferred regulatory interactions. Across multiple network constructions, simulations of single-gene perturbations consistently identified the Notch effector RBPJ as the key regulator predicted to shift MCC cells toward a more differentiated state. Experimental knockdown of RBPJ in an MCC cell line altered expression of differentiation-associated genes, reduced the expression of MCC markers, and drastically reduced cell growth. These findings identify RBPJ as a regulator of MCC lineage plasticity and candidate for targeted treatment, while highlighting the utility of probabilistic network modeling for prioritizing therapeutic targets in translational cancer research.

## Introduction

Neuroendocrine (NE) cancers are a heterogeneous class of tumors composed of cells with both neuronal and endocrine features and are often associated with neural lineage genes and hormone producing markers^1,2^. Among these, poorly differentiated NE carcinomas such as Merkel cell carcinoma (MCC) are especially aggressive and marked by high proliferation, drug resistance, and poor prognosis^3,4^. MCC is a neuroendocrine carcinoma of the skin and is often associated with Merkel cell polyomavirus (MCPyV) integration^5,6^. Histologically, MCC cells lack a clear pattern of differentiation, and signatures of their tissue of origin can be obfuscated^3,7,8^. This loss of lineage fidelity is often accompanied by alterations in pathways regulating cell state,^9^ such as TP53 and RB1^3^, which regulate stemness related genes including the Yamanaka factors^10,11^. The current standard of care for MCC is immunotherapy, which only about 50% of patients respond to, and among responders, only a fraction achieves a durable response^5,12^. Given that MCPyV is present in most MCC cases, immunosuppressed patients (who represent a highly vulnerable group for infection) are often ineligible for immunotherapy^5,6,12,13^. These challenges underscore the critical need for alternative therapies that target MCC-specific regulatory mechanisms.

Dedifferentiation in virus-positive MCC (VP-MCC) is driven by interactions between MCPyV small T antigen and L-Myc/MYCL, and large T antigen and Rb/RB1. This process is associated with changes in developmental potency, the ability of a cell to generate different cell types^11,14,15^. Totipotent and pluripotent stem cells can give rise to all or most cell types,^16–19^ whereas multipotent, oligopotent, and unipotent cells are progressively more restricted in what lineages they can form; differentiated cells represent the most specialized endpoint^17,19^. In cancer, cells can alter their potency and lineage to metastasize and to adapt to environmental pressures including immune responses and anticancer drugs^11,14,15,20–23^. For example, non-small cell lung cancers (NSCLC) treated with EGFR inhibitor can evolve into NE SCLCs post-treatment^24^. Conversely, reversing these changes may make tumors more sensitive to treatment^2,4,18,22,25^; in a murine model of treatment-resistant NE prostate cancer, knockout of the NE lineage-determining transcription factor (TF) ASCL1 was shown to restore sensitivity to androgen deprivation therapy^26^. Other examples have appeared in recent studies in lung^27,28^, breast^27,28^, and prostate cancers^11,26,29,30^, highlighting promising approaches to selectively target molecular mechanisms of differentiation and plasticity to make the tumors more vulnerable to existing therapies. However, the regulation of differentiation in MCC is still poorly characterized, restricting the development of such strategies.

The Notch signaling pathway is an important regulator of differentiation and plasticity in normal skin cells. Canonically, signaling is activated by the binding of a ligand (Delta-like or Jagged) to a transmembrane Notch receptor, which induces the cleavage and release of the Notch receptor’s intracellular domain (NICD). NICD then translocates to the nucleus where it forms a complex with recombinant signal binding protein for immunoglobulin kappa j region (RBPJ)^31–34^. This complex activates the expression of TFs involved in processes such as differentiation (HES and HEY)^35^ and cell cycle regulation (CCND1)^33^. When NICD is not present, RBPJ associates with transcriptional corepressors and acts as a repressor itself. Approximately 50% of virus-negative MCC tumors harbor mutations in the Notch pathway, including NOTCH1 and NOTCH2. Changes in the expression of Notch pathway members have been correlated with virus status, increased cell death, and prognosis^36^. However, there is no clear understanding of the influence of the Notch pathway on MCC cell growth and differentiation.

Here, we take a systems-based approach to integrate these concepts and identify mechanisms driving lineage plasticity and NE differentiation within MCC tumors. We analyzed single-cell RNA-seq data from twelve virus positive MCC tumors (six each from two independent datasets) to identify distinct subpopulations displaying varying levels of stemness and NE differentiation using CytoTRACE2^19^, a computational tool for predicting cellular developmental potency. We then identified key TFs that were differentially expressed between these subpopulations. Using these TFs, we defined a prior regulatory network and inferred additional regulatory relationships using ATAC-seq data. These networks were then pruned based on scRNA-seq data by the probabilistic Boolean network inference algorithm, BooleaBayes^28^. By modeling gene regulation probabilistically, BooleaBayes helps identify perturbations likely to shift system behavior. We determined which TF knockdowns would push MCC cells into a more differentiated and less stem-like state. Through our analysis, we identified several TFs with the potential to reverse MCC stemness, with RBPJ being the top-scoring candidate. We then characterized the effect of stable RBPJ knockdown in an MCC cell line and found that loss of RBPJ altered differentiation-associated gene expression and abrogated cell growth. Together, these findings support a role for RBPJ as a regulator of MCC stemness and reveal the potential of RBPJ inhibition as a therapy for MCC patients.

## Methods

### Single cell RNA-seq data processing

Our primary scRNA-seq dataset (GEO accession: GSE226438) consisted of eleven MCC tumors^37^. The data was processed using Seurat^38^, then subset to include only the six MCPyV positive (VP) tumors. The data was then filtered to cells with at least 500 transcripts, between 1,000 and 10,000 unique genes detected, and less than 5% mitochondrial genome. Genes found in less than 50 cells were removed. The data was then normalized using SCT normalization from the Seurat library and adjusted for cell cycle variation. The 2,000 most variable features were used to anchor and integrate the samples using SCT normalization. The secondary dataset (dbGaP accession no. phs002260) was processed using the same pipeline^39,40^.

### Clustering and dimensionality reduction

First, principal component analysis (PCA) was run on the data using the top variable features. The first thirty principal components (PCs) were used to construct a shared nearest-neighbor graph. Community detection was then performed on this graph using the Louvain algorithm with a resolution of 1.0, creating distinct clusters. Finally, uniform manifold approximation and projection (UMAP) embedding was calculated over the same 30 PCs. For the subclusters, a resolution of 0.25 was used to avoid over-clustering smaller populations^40^.

### Tumor cell classifier

To classify tumor versus non-tumor cells across our data sets, we trained a LASSO (least absolute shrinkage and selection operator) regression classifier on our primary data set and used it to predict the label of the cells in our secondary data set. For the training set, calculated tumor markers and cell-type signatures from ClusterMole^41^ were used to classify each cluster as tumor cells or immune/stromal cells. Then we trained the model using the most variable features, performed 10-fold cross validation to find an optimal value for λ, and trained an optimal model with this λ. LASSO regression was chosen to eliminate as many irrelevant features as possible. The optimal model contained 631 parameters and exhibited an accuracy of 0.933 on the test set. Once the model was defined, predictions were made for the cells of both datasets^40^.

### CytoTRACE2

Datasets were first filtered to only include tumor classified cells. Once filtered, PCA, SNN, and UMAP were carried out as above. Clustering was changed to a resolution of 0.4 to focus on biologically relevant communities. CytoTRACE2 was run on the SCT normalized tumor cell data, and absolute score, relative score, and predicted potency were added to the Seurat object^40^.

### Prior network construction

First, all differentially expressed genes were found between cells labeled by CytoTRACE2 as ‘Differentiated’ and ‘Multipotent’. Cells were grouped by potency, both positive and negative markers were included, a minimum of 10% of cells in either group had to express a gene for it to be considered, a minimum log fold-change of 0.25 was required, and the Wilcoxon Rank Sum test was used for statistical testing. Next, the set of differentially expressed genes was filtered to only include transcription factors with *p_adj_<0.05*. ARACNe was used to generate regulons^42,43^ for VP MCC and filtered to interactions where both the target and the source were in our list of potency-altered transcription factors. The DoRothEA network^44^ was filtered to interactions with at least a confidence of *C*, and then to edges where the source and target were in our list of transcription factors. Both lists of sources and targets were combined for each dataset, and the intersection of edges in both became our prior network^40^.

### ATAC-seq data processing

MCC cell-line (MKL1) ATAC-seq datasets (GEO accession no. GSE140505) were imported to R and converted to genomic ranges^45^. Intervals were restricted to standard hg38 chromosomes, ranges were trimmed to valid chromosome boundaries, and invalid peaks were removed. Peaks from all replicates were then merged into a nonredundant union peak set.

### ATAC-seq edge inference

Transcription factors from the prior network were extracted and used to retrieve corresponding DNA-binding motifs from CIS-BP^46^, JASPAR^47^, and HOCOMOCO^48^ motif libraries. For each TF with an available motif, FIMO^49^ was run against the union peak FASTA files to identify motif occurrences passing a threshold of *p<10^-4^*. Motif hits were filtered to those within ATAC peaks and later mapped to genomic motif-site coordinates. The sites were annotated to the nearest gene transcription start site (TSS) using a gene-level hg38 TSS reference, and only hits within 1 kb of a TSS were kept. These hits were collapsed to network edges with the TF as the source and the gene corresponding to the TSS as the target.

### Prior network edge filtering

First, for each transcription factor and its inferred edges, each edge was individually evaluated based on correlation of expression using BooleaBayes. Edges that passed a correlation threshold, *τ*, of 0.1 or 0.2 were added to the prior network. Additionally, if the source and target of any edges removed in the prior step remained, that edge was reintroduced to the network to account for TF co-regulation of specific genes. The full network was then run through the full BooleaBayes algorithm to identify the final network and determine regulatory rules.

### Transformation of expression values to probabilities

Single-cell expression data was log-transformed using ln(1+x) prior to transformation. Transformation was based on scBoolSeq^50^ classification, where for each gene, summary statistics describing its expression distribution was computed. Using these statistics, genes were labeled as bimodal, unimodal, or zero-inflated. Genes were labeled as bimodal if they showed evidence of multimodality based on dip test, Kurtosis, and bimodality index thresholds. Genes not meeting bimodal criteria were labeled zero-inflated if they showed both a low-density peak and high dropout rate, and all remaining genes were classified as unimodal. After classification, each gene’s expression was transformed according to its assigned category. Bimodal genes were modeled with a two-component Gaussian mixture model, and each cell was assigned a continuous values corresponding to the posterior probability of belonging to the high-expression component. Zero-inflated genes were binarized such that zero remained as 0, and non-zero values were set to 1. Unimodal genes were normalized only among non-zero expressing cells by rescaling values between upper and lower quantiles between the range of 0 and 1. This gene-specific normalization produced a continuous matrix of probabilities able to be used by BooleaBayes for rule fitting and other downstream analyses.

### BooleaBayes Processing

The prior network was defined as described above, and clusters were defined as the potency states ‘Differentiated’, ‘Unipotent’, ‘Oligopotent’, and ‘Multipotent’, with subclusters 0-5 for the differentiated cluster and 0-2 for the multipotent cluster to identify more cohesive clusters within these groups. The data scaled to probability values above was run using default parameters, and sinks and sources were kept, with a self-loop being added to the sources. Data were binarized for determination of binary cell states using gene-specific thresholds defined from the midpoint between the median probability values of two extreme phenotypic clusters (Differentiated_1 and Multipotent_1).

For gene *g*, the binarization rule was:

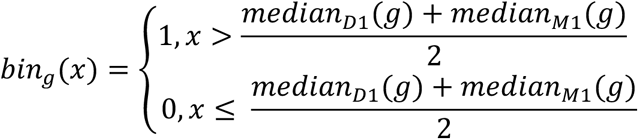

where *median*_*D*1_(*g*) *and median*_*M*1_(*g*) refer to the median probability of gene *g* being ON in clusters Differentiated_1 and Multipotent_1 respectively. The threshold, *τ*, for rules was set at 0.1 unless otherwise noted. That is, for a regulator *r*, where input *i* and *j* differ only by *r,* if:

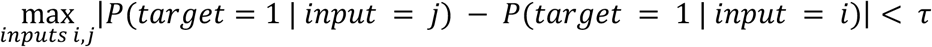

then *r* is deemed irrelevant and the edge is pruned. In other words, if changing the state of the regulator has a negligible effect on the predicted target output, the edge between them is removed^28^. For all applications of BooleaBayes, we exclusively used the RNA expression from the larger dataset (Das et al.) to reduce the impact of weakly supported low-confidence regulatory relationships.

### Pseudo data generation

Randomized synthetic RNA-seq data was generated to test the predictive capacity of this workflow as a negative control. The log-transformed scRNA-seq data from the primary data set was read, keeping the gene and cell labels as well as determining the global minimum and maximum expression values. For each gene-cell entry, a value was generated uniformly at random from the observed expression range. This maintained the original structure and bounds while removing any real biological signal or correlations.

### Dynamical importance calculation

Dynamical importance is a quantification of how much the removal of a certain node changes the largest eigenvalue (*λ*) of the network adjacency matrix, which governs the behavior of many dynamical processes on a graph. The right and left eigenvectors of the adjacency matrix, *A*, are denoted by *u* and *v*, such that *Au* = *λu* and *v*.*A* = *λv*.. To avoid the cost of recomputing *λ* for the ablation of each element, the dynamical importance of a node *k* can be estimated to first order as^51^:

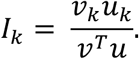

### Cell culture and chemicals

Characteristics of HDFn^52,53^ cells and Merkel cell carcinoma cell line WaGa^39,45,54,55^ have been previously reported. The fibroblast cell line HDFn was obtained from ATCC (PCS-201-010). All the cell lines were cultured at 37°C under 5% CO_2_. HDFn cells were cultured in complete DMEM medium (Corning, cat 10-013-CV), containing 10% FBS, 1% GlutaMAX (GIBCO), 10 U/mL pen/strep (GIBCO cat 35050061), and 10 mg/mL streptomycin (GIBCO). WaGa cells were cultured in RPMI-1640 medium (Corning, cat 10-040-CV) containing 10% FBS, 1% GlutaMAX (GIBCO), 10 U/mL pen/strep (GIBCO cat 35050061). 293T cells were cultured in Dulbecco’s modified Eagle medium (DMEM; Corning, cat 10-013-CV)) supplemented with 1% Pen/Strep (GIBCO cat 35050061), 1% GlutaMAX, and 10% FBS.

### Transformation of Competent Cells

Competent DH5α *E. coli* cells were transformed with 2-5 μl of the shRBPJ plasmid or SHC002 using the protocol supplied with competent cells. DH5α cells were removed from the -80 freezer, and 50 μl of cells were added to a 2.5 ul DNA Eppendorf containing the vector. The mixture was incubated on ice for 30 minutes. Agar plates were transferred from the cold room into the 37°C incubator, and the water bath was set to 42 °C. The bacteria were heat shocked in the 42°C water bath for 60 s then immediately returned to ice for 2-5 minutes. Afterwards, 100 μl of LB broth was added to the cell suspension. The tubes were gently mixed by tapping, the suspension was spread onto LB plates with ampicillin, and the plates were incubated at 37° for 16 h. A single colony was picked and cultured in 2 ml LB media with 100 μg/mL ampicillin and incubated at 37°C with gentle shaking for 16 h. This 2 ml culture was added to 100 ml LB media containing ampicillin and grown at 37°C with gentle shaking at 250 rpm for 16 h. Plasmid vectors were extracted using the ZymoPURE II Plasmid Midiprep Kit (D402).

### Production of shRNA lentivirus

HEK293 cells were used to produce lentivirus from the lentiviral vector and packaging plasmids. The lentiviral packaging plasmid psPAX2 and envelope plasmid pMD2.G were gifts from Didier Trono (Addgene plasmids 12260 and 12259). Transfections were performed using Lipofectamine 3000 (Invitrogen, #L3000001). Approximately 24 h before transfection, 3 × 10^6^ HEK293 cells were seeded into 10 cm tissue culture plates in 10 mL growth medium without antibiotics and incubated overnight at 37°C in 5% CO2. Cells were approximately 70% confluent at the time of transfection. Sixteen hours after transfection, the transfection medium was replaced with 10 mL fresh complete growth medium, and cells were incubated at 37°C for an additional 24 h. Lentiviral supernatants were harvested 48 h after the start of transfection and filtered through a 0.45 μm low-protein-binding filter to remove cellular debris.

### Transducing target cells with shRNA lentivirus

WaGa cells were plated at 1 × 10^7^ cells in a 10 cm tissue culture dish in 5 mL complete growth medium and incubated at 37°C in a humidified atmosphere containing 5% CO2 prior to infection. WaGa cells were infected the same day with lentivirus. Cells were treated with 5 mL of shRBPJ virus (TRCN0000016203, TRCN0000016204, and TRCN0000016205) or 5 mL of SHC002 virus together with polybrene at 2 μg/mL. Viral supernatant was added to the cells, and transduction proceeded for 48 h. The virus-containing medium was then removed and replaced with fresh growth medium. Cells were incubated for an additional 24 h to allow the shRNA to reach maximum effect. Because the resistance gene conferred puromycin resistance, puromycin was added at 1 μg/mL for 2–3 days until resistant colonies could be identified.

### Quantitative real-time PCR

Cellular RNA was extracted using the Quick-RNA Miniprep Kit (Zymo, cat. 15596026) according to the manufacturer’s instructions. RNA samples were reverse transcribed into cDNA using the High-Capacity cDNA Reverse Transcription Kit (Thermo Fisher Scientific, cat. 4368814). Real-time PCR was performed using the Applied Biosystems StepOne system with SYBR Green RT-PCR Master Mixes (Thermo Fisher Scientific, cat. A25742). PrimePCR SYBR Green assay primers (see Table 1) were used to amplify genes of interest and the housekeeping gene. Data were acquired as threshold cycle (Ct) values. ΔCt values were determined by subtracting the average housekeeping gene Ct value from the average target gene Ct value. Because amplification efficiency of the target genes and internal control gene was assumed to be equal, relative gene expression was calculated using the 2^−ΔΔCt method. Each measurement was performed in triplicate and repeated three times.

**Table 1.**
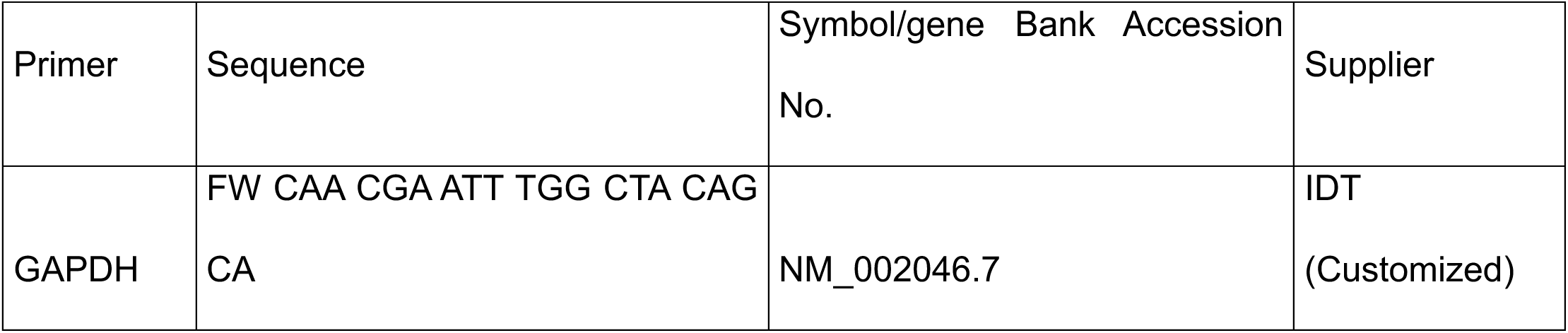
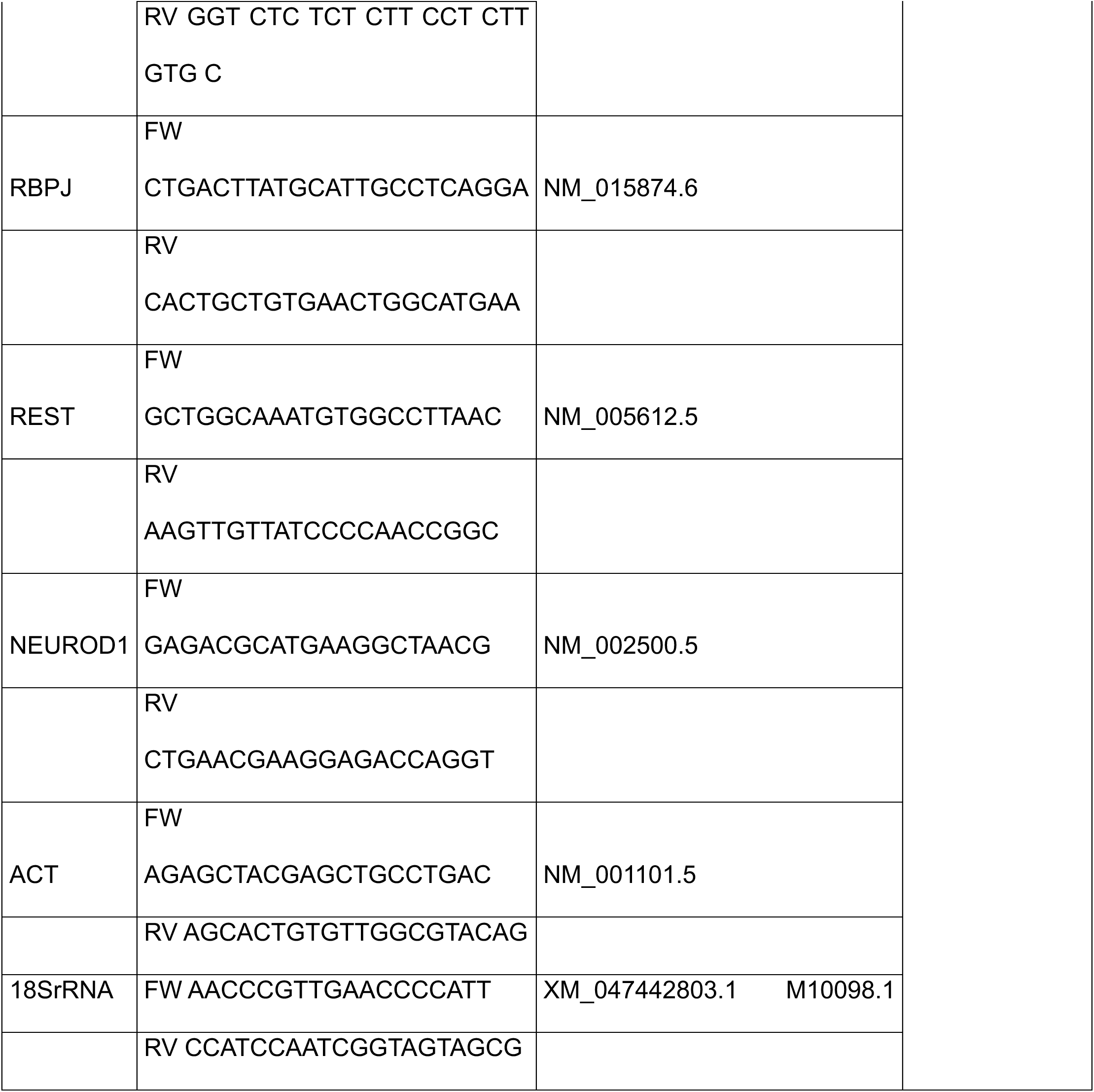
Primers used in RT-qPCR.

### Western Blot

Following puromycin selection, WaGa cells were seeded in 10 cm^2^ tissue culture plates. Cells were washed with cold PBS and lysed in RIPA buffer (Thermo Fisher Scientific, cat. 89901) supplemented with EDTA-free protease inhibitor cocktail (Roche, cat. 04693132001). Lysates were centrifuged for 10 min at 32,310 g at 4°C, and the supernatant was collected. Protein concentration was determined using the Pierce BCA Protein Assay Kit (Thermo Fisher Scientific, cat. 23227). Protein samples were denatured in 4× Laemmli sample buffer containing 10% β-mercaptoethanol (Sigma-Aldrich, cat. M6250) by boiling at 95°C for 10 min. Samples were then loaded onto SDS-PAGE gels and electrophoresed for 80 min at 120 V. Proteins were transferred to nitrocellulose membranes using a semi-dry transfer method at 1.5 A for 30 min. Membranes were blocked and incubated with primary secondary antibodies according to manufacturer recommended conditions. Anti-RBPJ antibody was used to detect RBPJ, and anti-Vinculin antibody was used as a loading control.

**Table 2.**
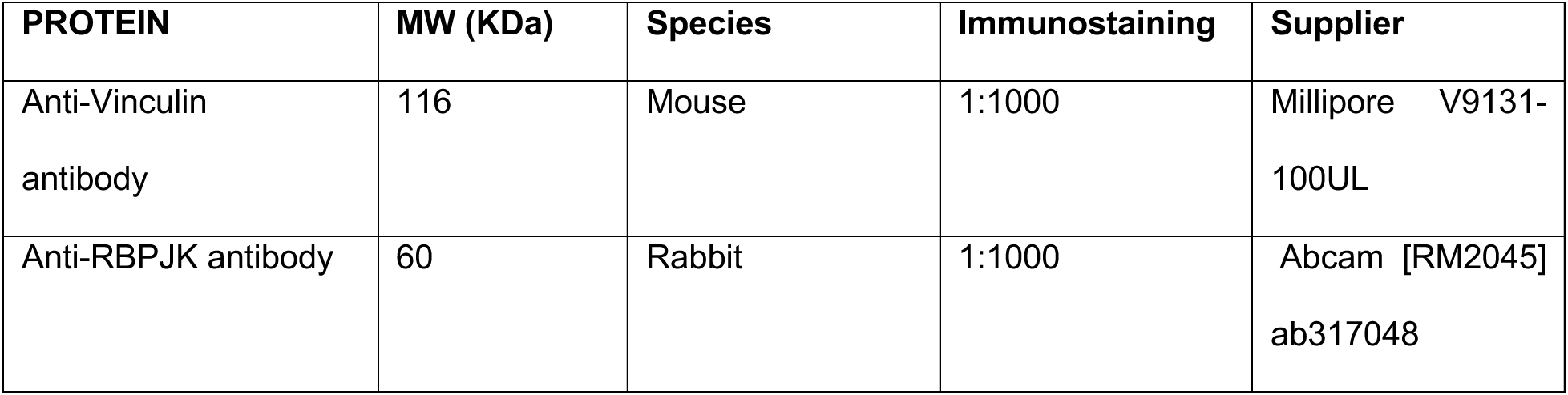
Primers used in RT-qPCR.

| PROTEIN | MW (KDa) | Species | Immunostaining | Supplier |
| --- | --- | --- | --- | --- |
| Anti-Vinculin antibody | 116 | Mouse | 1:1000 | Millipore V9131-100UL |
| Anti-RBPJK antibody | 60 | Rabbit | 1:1000 | Abcam [RM2045] ab317048 |

### Cell Viability and Cell Proliferation

WaGa cells were seeded in 96-well plates at a density of 20 × 10^4 cells in 100 μL. Water-soluble MTT (thiazolyl blue tetrazolium bromide; Sigma-Aldrich, cat. #M5655) was dissolved in DPBS to 5 mg/mL. At each measurement time point, 10 μL of 5 mg/mL MTT was added to each 100 μL cell suspension, and plates were incubated at 37°C for 2 h. After incubation, 100 μL of DMSO/30% SDS was added to each well. Plates were then wrapped in foil and shaken on an orbital shaker for 10 min at 37°C. Absorbance was read at 540 nm using a BioTek Synergy LX plate reader. HDFn cells were seeded in 96-well plates at the same density of 20 × 10^4^ cells in 100 μL and processed similarly. For HDFn cells, after incubation with MTT, 100 μL warmed DMSO was added to each well, and plates were shaken on an orbital shaker at 400 rpm for 10 min at 37°C before reading at 540 nm on the BioTek Synergy LX plate reader.

### Bulk RNA-seq

WaGa cells were transduced with lentiviral vectors containing shRBPJ or SHC002 sequence as a control. Cells were selected with puromycin for 48 h and harvested in triplicate. RNA was purified using the Quick-RNA Miniprep Kit (Zymo, cat. 15596026). Isolated RNA was subjected to quality control using an Agilent 2100 Bioanalyzer, and all samples passed quality control. RNA samples were then subjected to library preparation and sequencing on the Illumina NovaSeq 6000 platform by Novogene.

Bulk RNA-seq count data were imported from a count matrix. Cell line and condition were labeled for each sample. Gene identifiers were kept as Ensembl IDs with annotated gene symbols. Prior to differential expression analysis, the genes were filtered to only keep those with at least 10 counts in at least 3 samples. Differential expression analysis was performed in DESeq2^56^. Gene Ontology (GO) term enrichment analysis was performed on significantly (p<0.05 and abs(log_2_(fold-change))>0.5) upregulated and downregulated genes to identify biological processes associated with each expression pattern. Gene set enrichment analysis (GSEA) was used to evaluate changes across ranked gene expression profiles without the requirement of a significance threshold. To compare bulk RNA-seq results with the scRNA-seq analysis, differentially expressed markers related specifically to cell states were identified within the scRNA-seq data using the FindMarkers function in Seurat and then visualized in the bulk RNA-seq data.

*SCENIC*

The Single-Cell Regulatory Network Inference and Clustering (SCENIC) tool was used to infer gene regulatory networks with the pySCENIC version 0.12.1 docker image. The standard pipeline was followed, using GRNBoost2 to infer gene regulatory networks, followed by regulon prediction with cisTarget database, and single-cell transcription factor activity estimation with AUCell.

## Results

### MCC tumors contain subpopulations with distinct differentiation states

To better understand the poorly differentiated phenotype of MCC we first sought to characterize the heterogeneity in developmental potency of the tumor cells. Following classification and selection of tumor cells (Figure 1, B and C) in two independent scRNA-seq datasets, we used the CytoTRACE2 deep learning framework to calculate absolute potency scores for each cell (Figure 1, D and E). This revealed populations of cells classified as Differentiated, Unipotent, Oligopotent, and Multipotent in both datasets. In our first dataset, the percentage of cells in each category was 45%, 21%, 27%, and 7% respectively. In the second dataset, the percentages were 23%, 56%, 16%, and 5%. This corresponds with our expectation of poorly differentiated tumors, as the majority of cells in both datasets were classified with a higher potency than differentiated^54^. Differences in proportions could be due to expected variation between datasets, but it is worth noting that one of the patients form the first dataset had already begun immunotherapy via pembrolizumab treatment when the sample was collected, which may impact the potency spectrum^57^. In contrast, a similar analysis performed on pancreatic NE tumors, which are morphologically well-differentiated, revealed only differentiated cells and almost no cells classified with higher potency (personal communication, M. Padi and E. Mayere).

**Figure 1.**
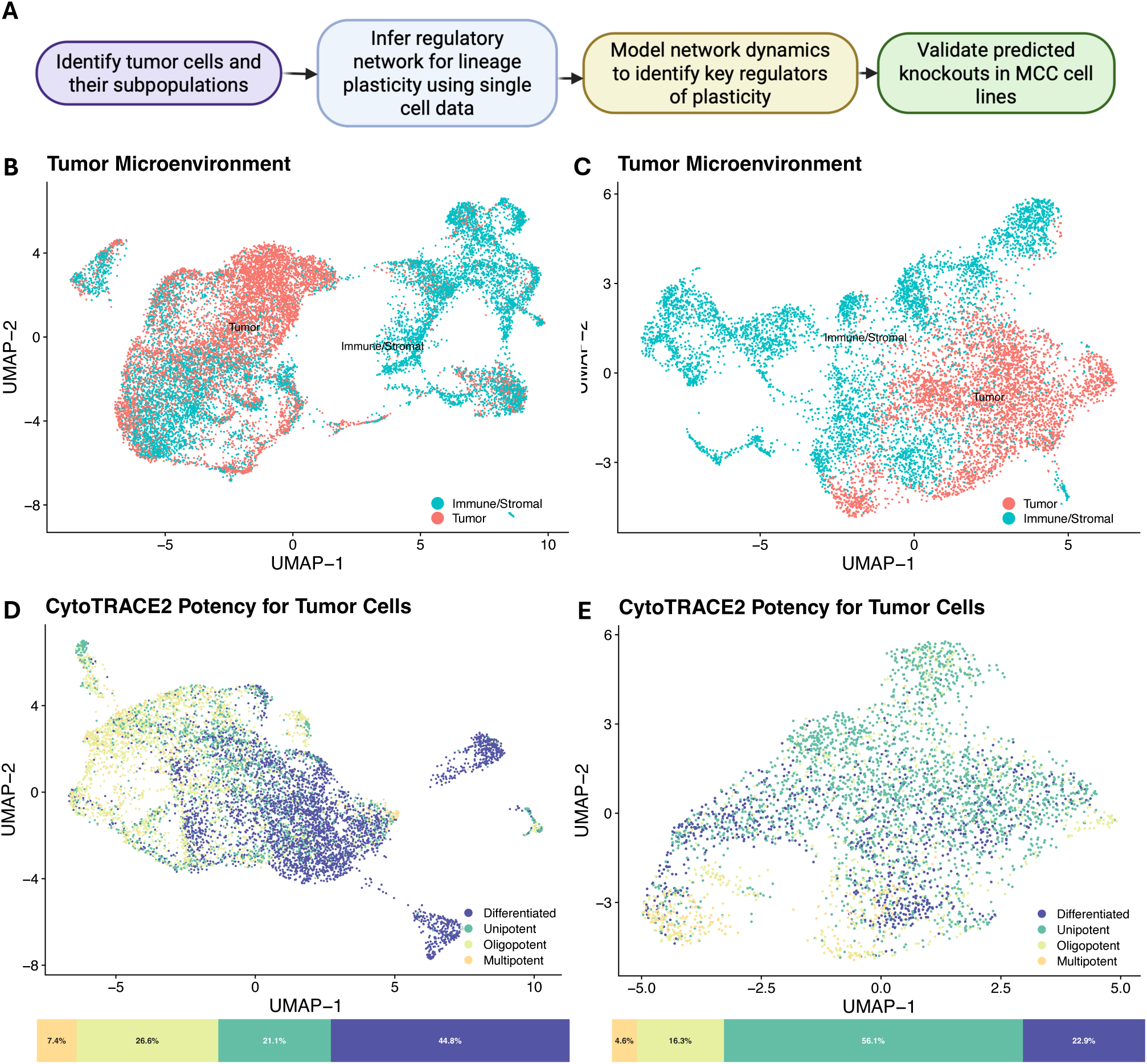
Distinct levels of differentiation exist within MCC tumors. **A**. General workflow used for this project. **B,C**. UMAP colored by tumor classification from LASSO classifier prediction. **D,E**. UMAP filtered to only tumor cells and colored based on potency calculated by CytoTRACE2. Below the UMAP, bar plots display the distribution of potency within tumor cells for each dataset. Differentiated and unipotent cells are the majority, but there do exist smaller oligopotent and multipotent populations in both independent datasets. B,D. Dataset from Das et al. C,E. Dataset from Frost et al.

### Regulatory network construction using improved approaches for data normalization and binarization

To model the regulatory network dynamics governing the transition from multipotent to differentiated cell states, we used Boolean networks. Boolean networks are directed graphs where the nodes are elements and the edges represent how those elements regulate each other^58^. For a gene regulatory network, the nodes are TFs and the edges represent activation or inhibition by a regulator to its target. Each node is in a binary state of ON (1) or OFF (0) and has an associated Boolean rule or logic function that determines its state based on input nodes using AND, OR, and NOT operators. The dynamics of the network can then be simulated using the set of rules for all of the nodes in the network, by setting an initial state for all nodes and iteratively updating the node states according to their rules. Since there is a limit to the number of possible states of a network, the system eventually converges to a steady state, representing a cellular phenotype.

To create a Boolean network that can model the transition between multipotency and differentiation, we first constructed a prior network using differentially expressed (DE) transcription factors between Differentiated and Multipotent cells (Figure 2B). This network contained 61 nodes and 100 edges derived from ARACNe, which uses mutual information to identify MCC-specific direct associations between TFs and target genes, and DoRothEA, which curates regulatory relationships from multiple contexts based on experimental data like ChIP-seq. ARACNe and DoRothEA represent the state-of-the-art methods for constructing high-confidence regulatory networks from MCC RNA-seq data and manually curated databases, respectively. These methods have low sensitivity but focus on the most reliable regulatory relationships, making them a good starting point for modeling network dynamics.

**Figure 2.**
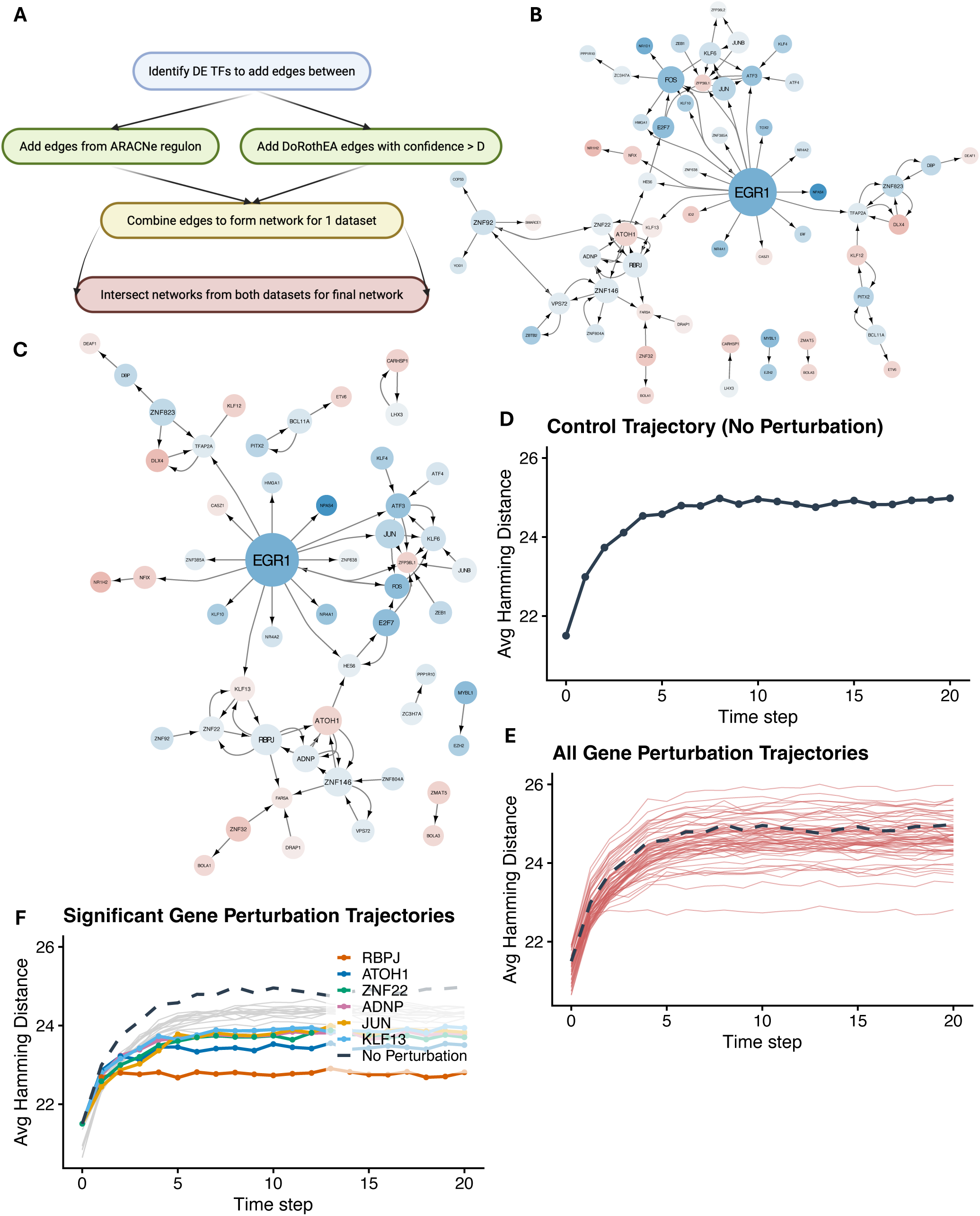
Prior network allows for predictions of key regulators. **A**. Workflow of prior network construction **B**. Prior network contrasting multipotent versus differentiated states: 61 nodes and 100 edges sourced form DoRothEA and ARACNe interactions. Node size scales with outdegree of the node. Color indicates differential expression of the node between differentiated and multipotent cells, where red is upregulated in differentiated cells and blue is downregulated. **C**. Network after BooleaBayes pruning: 52 nodes 77 edges. **D**. Multipotent cells unperturbed: average Hamming distance of multipotent cluster 1 cells to the average state of differentiated cluster 1 cells over 20 time steps and 10 simulations. **E**. Single gene OFF perturbation: average Hamming distance from differentiated state over time. Dashed line is the control trajectory. **F**. Trajectories over time of genes that significantly lower distance to differentiated state, with the 6 most significant annotated.

To fit the network to MCC tumor data, we used BooleaBayes, a probabilistic Boolean network inference tool that generates regulatory rules based on scRNA-seq profiles. Rules for a gene are calculated by estimating the probability that the gene is ON under different combinations of regulator states from the data, which accounts for biological heterogeneity and noise, making it particularly useful for studying heterogeneous cancer cell states^28,40^. To infer these rules, the expression data must be transformed to the range [0,1] such that each value represents the probability of that gene being ON in that cell. The details of this transformation and the subsequent binarization can have nontrivial consequences on the final model behavior. We first tried several default binarization methods from the BooleaBayes package, including mapping non-zero values to 1 and zero to 0, using a Gaussian mixture model (GMM), and quantile thresholding, but none of these methods was able to capture the difference between multipotent and differentiated cells following binarization. Thus, we implemented a combined approach based on the package scBoolSeq using log1p-transformed data instead, where we first categorized the distribution of the gene expression into one of three groups: Unimodal, Bimodal, or Zero-inflated (Figure 3C). These categories were then normalized using quantile thresholding, GMM-based fitting, and threshold-based binarization, respectively, to scale gene expression values to the range [0,1]. Binarization for Boolean network simulations was then performed by setting the threshold as the midpoint between the median probability value for the multipotent and differentiated cell clusters. This binarization strategy preserved the patterns of differential expression observed in the original scRNA-seq analysis and produced more biologically realistic results. For example, for all source nodes (nodes that are only regulated by themselves), rules were inferred by BooleaBayes to be logically consistent, i.e. if gene A activates itself, and gene A is ON, then gene A should remain ON despite the probabilistic nature of the rules. Since the regulator state and the target state of a self-loop are read from the same expression value, they should be binarized to the same value in every cell, so a node that is ON remains ON, and one that is OFF remains OFF, which this strategy captures across different expression distributions. Following rule fitting and edge pruning by BooleaBayes, the network contained 52 nodes and 77 edges (Figure 2C).

**Figure 3.**
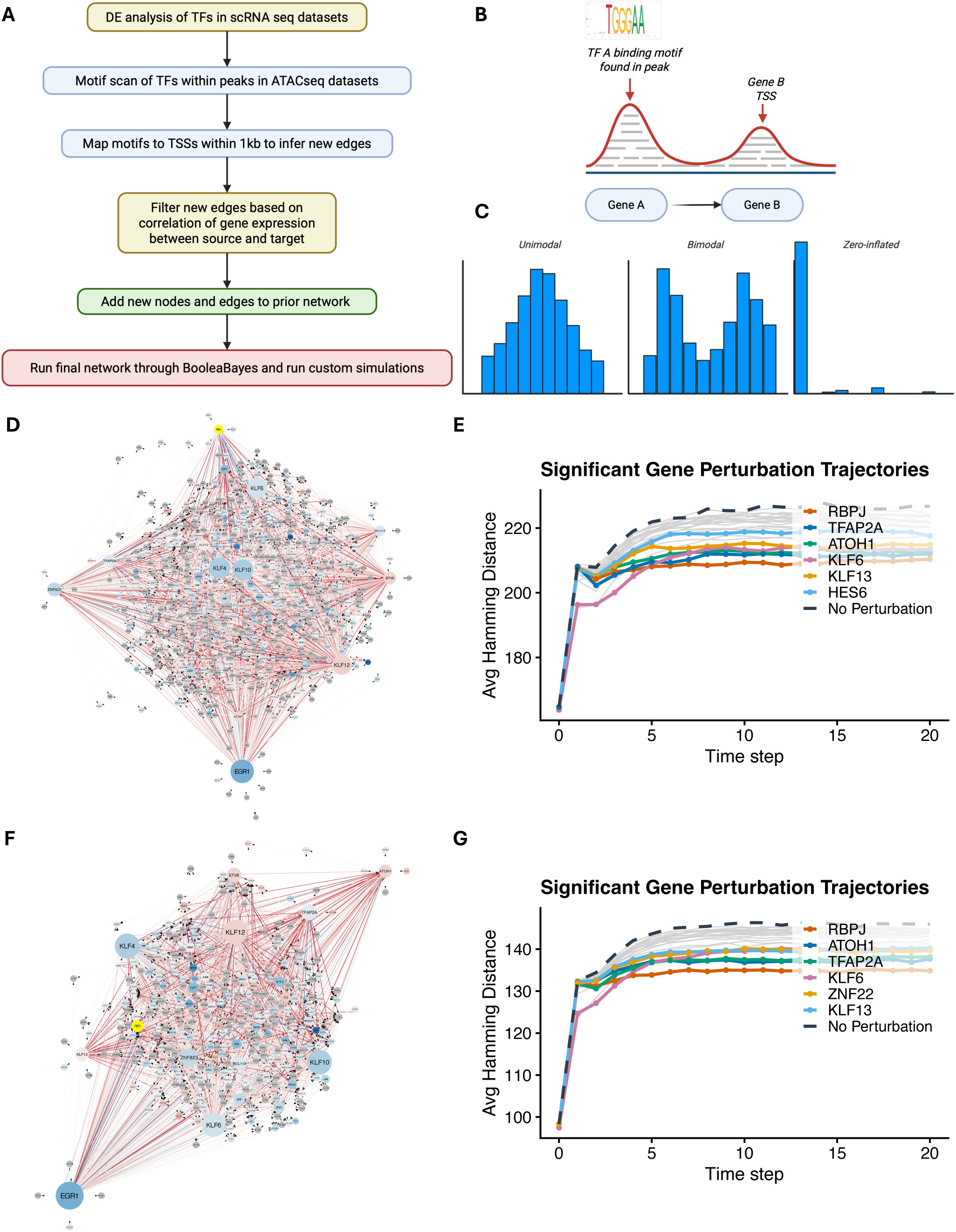
New workflow to infer edges from scRNA-seq and ATAC-seq data. **A**. Workflow of our network enrichment to provide more system specific and data driven edges to the prior network. **B**. Diagram illustrating how edges are added from the ATAC peaks. **C**. Visualization of classes for gene normalization; each gene is scaled 0 to 1 based on its classification. **D-G**. Enriched networks maintain RBPJ as the major regulator of differentiation. **D**. Enhanced network A with *τ* = 0.1: 626 nodes and 4459 edges. Size corresponds to outdegree, and color to differential gene expression between differentiated and multipotent cells. RBPJ is highlighted in yellow. Color of the edges represents signed strength of interaction where red is a larger positive magnitude, blue a larger negative magnitude, and white represents smaller values. **F**. Enhanced network B with *τ* = 0.2: 316 nodes and 2402 edges. **E and G**. Trajectories over time of genes that significantly lower distance to differentiated state.

### Network simulation predicts RBPJ as a key regulator

It has previously been demonstrated that forcing de-differentiated cells towards terminal differentiation can make them more responsive to treatment^4,11,25^. To identify perturbations that would push a multipotent cell to a more differentiated state, we simulated the trajectory of each cell in our perturbation cluster following the ‘knockdown’ of a single gene by keeping it in the OFF state for the simulation and stochastically updating the state of all other nodes according to their rules over 20 iterations or ‘time steps’. This was done 10 times for each cell in the perturbation cluster for each gene, and we calculated the average Hamming distance between the perturbed cells and the average state of the reference cluster for each time step.

Both the Multipotent and Differentiated clusters were broadly distributed in transcriptional UMAP space (Figure 1C), suggesting that an average computed across all cells within either group would not create a representative cell state. However, each population contained more tightly clustered subpopulations, whose average states were more likely to reflect coherent regulatory programs. We therefore reclustered the Multipotent and Differentiated cells separately and selected Multipotent_1 as the “perturbation” cluster and Differentiated_1 as the “reference” cluster for downstream simulations.

As a control, we first performed 10 simulations of the cells in the perturbation cluster evolving over 20 time steps with no perturbations (Figure 2D). The cells reached a steady state with an average Hamming distance of ∼25 from the reference cluster. Then, each TF was fixed to OFF and simulated as described above (Figure 2E). We then selected all TFs whose final Hamming distance was significantly lower than the average final distance over all knockouts (Figure 2F; Mann-Whitney U test). 20 TFs were found to push the cells significantly closer to the reference cluster; many of these TFs have previously reported roles in regulation of differentiation. Knockout of the Notch effector RBPJ caused the greatest shift in Hamming distance toward the reference cluster, followed by the lineage-determining factor of Merkel cells, ATOH1.

While the average final Hamming distance across all cells in the perturbation cluster may identify important perturbations, it does not capture the response of single cells and cannot characterize the statistical significance of this shift. To address this, we analyzed the distribution of the Hamming distance for the final two time-steps for each gene and the control. Though all the distributions were unimodal, the most significant genes had higher densities below the average of the control (Supplemental Figure 1). We then determined which perturbations have a significantly higher proportion of cells below the mean value of the control compared to the control using the two-proportion z-test.

RBPJ was the most significant with roughly 68% of the cells settling to a state closer to the reference differentiated state than the control (*p*=1.36e-87). Therefore, we hypothesized that RBPJ is involved in the regulation of lineage differentiation in MCC and that knocking down RBPJ could lead to a more differentiated phenotype in MCC cells.

### Enrichment of prior network with ATAC-seq inferred edges

Published data on regulatory interactions specific to MCC is sparse, and as such, the prior network is thought to be largely incomplete. As a rare cancer, the mechanisms driving MCC involve unusual regulators like ATOH1 and L-Myc and may not be well-represented in databases like DoRothEA. ARACNe provides a small set of MCC-specific regulatory interactions but does not have the sensitivity to capture the majority of target genes. Thus, we sought to improve our network by generating regulatory edges based on TF motif occurrence and chromatin accessibility using ATAC-seq data from the MKL1 MCC cell line (Figure 3, A and B).

Through this workflow, we were able to add TFs and their regulatory relationships by identifying TF binding motifs that are accessible, mapping them to proximal TSSs, and keeping only those where expression of the regulator and the target are correlated with coefficient, *τ*, greater than 0.1, meaning only biologically feasible edges. We used this workflow to generate a new larger network (A) with 626 nodes and 4459 edges (Figure 3D). To check the robustness and sensitivity of our network models, we also generated a sparser network (B) with a *τ* of 0.2 (See Methods) with 316 nodes and 2402 edges (Figure 3F). Binding and Expression Target Analysis (BETA) on the ST-MYCL-EP400 complex identified 103 direct TF targets for L-Myc in MCC, suggesting that networks of this scale are reasonable from a biological standpoint^59^. However, as a computational model, network A was too large to be practical for downstream inference and simulation, and even network B approached the upper limits of feasibility in terms of both computational overhead and interpretability. Even so, modeling of perturbations in both networks A and B supported RBPJ as the most significant perturbation compared to the control (Figure 3, E and G).

### Network structure supports importance of RBPJ

To investigate why RBPJ was consistently selected as the top TF, we used randomized pseudo data in our workflow to determine if the behavior of RBPJ is driven by the tumor scRNA-seq data or only by the prior network structure. Using the small BooleaBayes network (Figure 2A) and its inferred rules from randomized data, we simulated single gene perturbations on the same multipotent cells and found that none led the system closer to the average differentiated state compared to the control (Figure 4, A and B), demonstrating that the rules inferred from MCC-specific scRNA-seq data drives the prediction of RBPJ. We next looked at dynamical importance, a measure of how much each node contributes to the network dynamics compared to its ablation. For the prior network, we found that RBPJ had the highest dynamical importance and that dynamical importance > 0 was a strong predictor of significant perturbations and not correlated to outdegree (Figure 4, C and E). When applied to the larger 626 node network A, EGR1 had the highest dynamical importance which was more closely correlated to outdegree (Figure 4, D and F). To investigate whether this was just due to the addition of many ‘sink’ nodes, which were 94% of the nodes in network A, we removed them to look at the core network and found that despite being comparable in number of nodes, network A had many more edges in this subset than the prior network (Figure 4, G and H). In summary, network structure by itself is not sufficient to predict master reguators of stemness. Despite not exhibiting high dynamical importancde, the RBPJ knockout leads to the greatest number of changes aligned with a transition to the differentiated state, highlighting the importance of the Boolean rules for fully capturing network dynamics.

**Figure 4.**
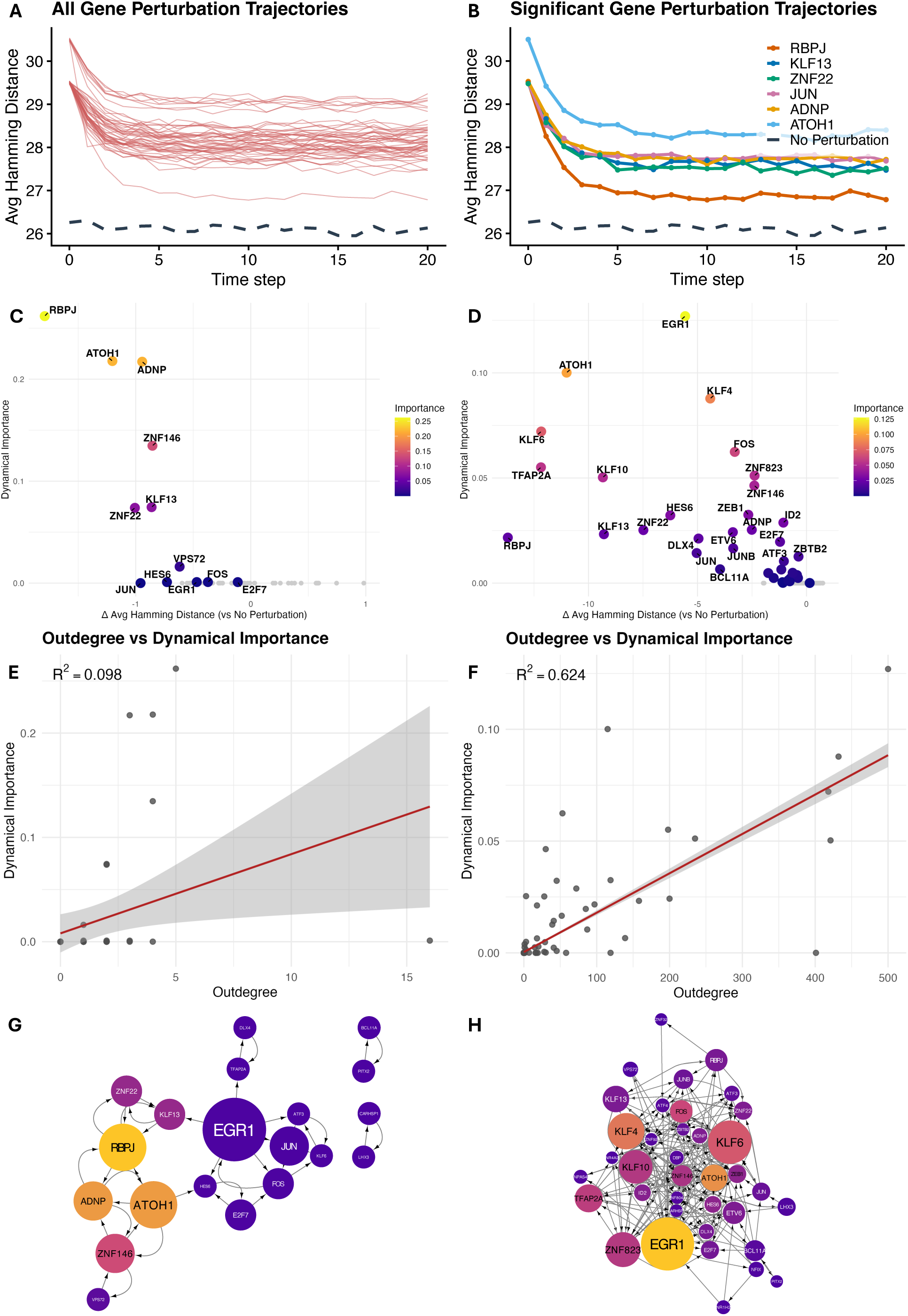
Network structure highlights importance of RBPJ and other TFs. **A**. Single-gene OFF perturbation on randomized data using original 52 node network. Average hamming distance between differentiated and multipotent states over 20 time steps; dashed line is the control trajectory. **B**. Trajectories for the 6 most significant genes in 2F under randomization. **C, D**. Dynamical importance versus perturbation significance (change in average Hamming distance to the differentiated state relative to control); more negative indicates a greater shift toward the differentiated state, colored by dynamical importance, for the 52 node network (C), and the enriched 626 node network (D). **E,F.** Dynamical importance versus outdegree with a linear least squares fit (red), 95% CI, and R2, for the same 52 and 626 node networks respectively. **G,H**. Core networks where all sinks and sources were removed for prior network (G) and network A (H). Size corresponds to outdegree and color represents dynamical importance as in C,D.

### Knockdown of RBPJ disrupts neuronal gene expression and proliferation in WaGa cells

We next sought to characterize the effect of silencing RBPJ in an MCC cell line. We transduced WaGa cells with shRBPJ or a control scramble non-targeting shRNA (shControl) on two independent occasions to produce biological replicates. Knockdown of RBPJ was confirmed with RT-qPCR, where we saw a reduction of RBPJ by ∼75% relative to the control in Biological Replicate 1 (BR1) and by ∼59% in Biological Replicate 2 (BR2) (Figure 5, A and B). We also measured expression of NEUROD1, a known regulator of neural differentiation. NEUROD1 was overexpressed following RBPJ knockdown in both BR1 and BR2 by 68% and 27% respectively (Figure 5, C and D). Western blot results confirmed decrease of RBPJ protein levels in shRBPJ cells compared to the control (Figure 5E). Cells expressing shRBPJ grew slower than cells expressing shControl. To quantify this and compare the effect of RBPJ on cell growth in cancer and normal cells, we transduced HDFn cells (neonatal human dermal fibroblasts) with shRBPJ or shControl and performed an MTT assay over 96 hours for the HDFn cells and over 168 hours for WaGa cells. The shorter time scale for HDFn was chosen so that these slow-growing primary cells were not maintained in the same growth media for an excessive time period. Viability metrics were normalized relative to time 0. In HDFn cells, both cell lines showed approximately linear growth over the 96-hour period, with shRBPJ cells increasing by ∼20% of Time 0 per day and shControl cells increasing by ∼12% of Time 0 per day (Figure 5F). By the end of the 96 hours, control HDFn cells reached 1.49 times the cells present at time 0, whereas shRBPJ expressing HDFn cells reached 1.87 times the Time 0 value. This is consistent with previous reports showing that silencing of RBPJ can increase proliferation in fibroblasts^60,61^.

**Figure 5.**
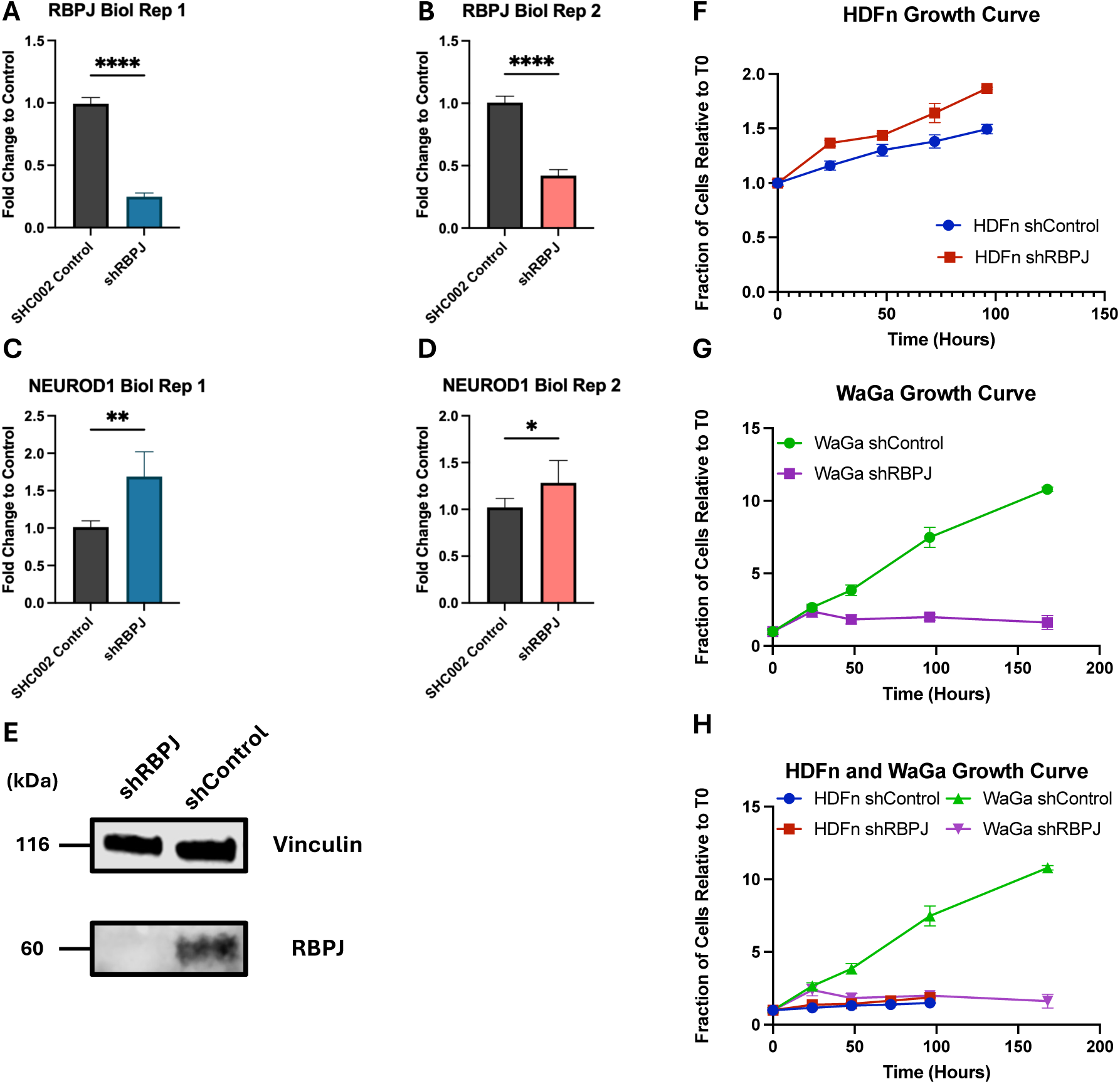
shRBPJ disrupts neuronal gene expression and proliferation in WaGa cells. WaGa cells and HDFn (neonatal human dermal fibroblasts) were transduced with shRBPJ. **A-D.** Relative mRNA levels of RBPJ and NEUROD1 was measured by RT-qPCR in WaGa cells across two biological replicates. Statistical significance was determined by unpaired 2-sample, 2-tailed t test. (****p<0.0001;**p<0.005, *p<0.05). **E**. RBPJ protein levels were assayed in WaGa cells with shRBPJ or non-targeting control (shControl) by immunoblotting **F-H**. Relative number of cells over time in WaGa and HDFn cells measured by MTT assay.

In WaGa cells, both cell lines initially increased in number for the first 24 hours, with the shControl reaching 2.68 times Time 0 and the shRBPJ reaching 2.38 times the cells at time 0. After this point, the control WaGa cells continued to grow at a rate of 138% per day, whereas the shRBPJ transduced cells decreased by 9.7% per day (Figure 5G). By 168 hours, control WaGa cells reached 10.80 times the number of cells of Time 0, while RBPJ silenced WaGa cells had fallen to 1.62 times the starting value. This demonstrates that silencing of RBPJ markedly reduces the growth of WaGa cells, opposite to the effect observed in non-cancerous HDFn skin cells. Beginning at 24 hours, shRBPJ expressing WaGa cells decreased in number, and by 96 hours their total growth was comparable to that of the much slower growing HDFn cells. Together, these findings support the hypothesis that in MCC cells, RBPJ is involved in the regulation of NE differentiation associated genes and is required for sustained proliferation.

### RBPJ silencing causes wide ranging transcriptional changes in WaGa cells

To better characterize the transcriptional response to RBPJ knockdown, we performed RNA-seq on two sets of independently transduced shRBPJ and shControl WaGa cells (i.e. 2 biological replicates), with three samples from each cell line (3 technical replicates). Principal Component Analysis showed samples clustered by both biological replicate and condition along PC1 and PC2, which accounted for 72% and 13% of the variance respectively (Figure 6A). The PCA shows that, despite the cell lines transduced at different times clustering separately from each other, the RBPJ knockdown had a similar effect on both biological replicate cell lines, underscoring the robust impact of this perturbation. We then visualized the most differentially expressed genes that were reproducible between the biological replicates (Figure 6, B and C). RBPJ was downregulated by 71% in shRBPJ expressing cells relative to the control, confirming that the knockdown was effective. Among the most significantly DE genes, Cannabinoid receptor 1 (CNR1), a regulator of neural progenitor differentiation, is strongly downregulated in the shRBPJ condition^62^. In contrast, Yamanaka factor MYC, as well as DUXA and its targets LEUTX and ZSCAN4, which together modulate zygotic genome activation (ZGA), were upregulated^63^. These transcriptional changes are consistent with dysregulation of MCC markers in shRBPJ transduced cells.

**Figure 6.**
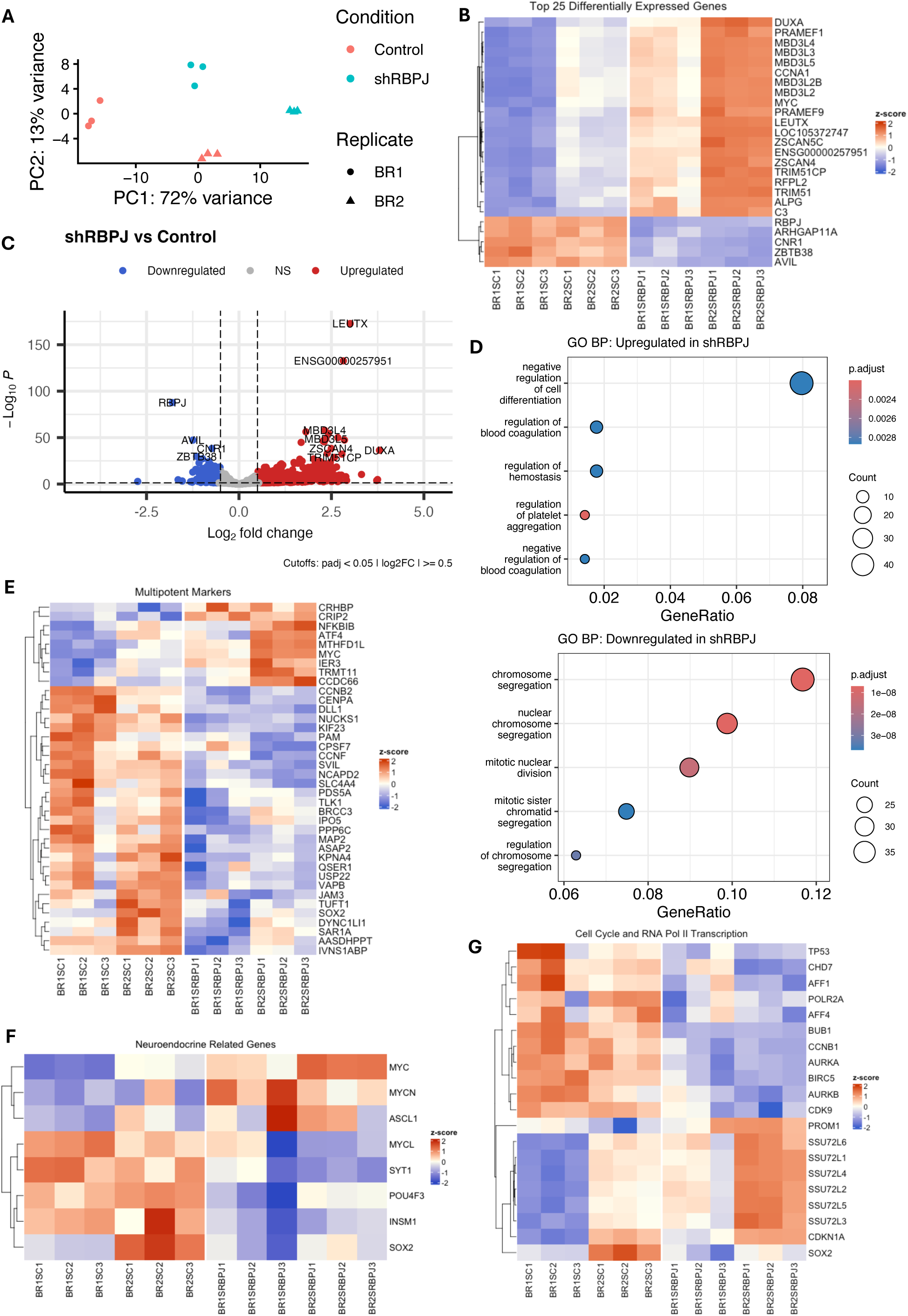
Bulk RNA-seq reveals broad transcriptional changes following RBPJ knockdown in WaGa cells. RNA-seq was performed on WaGa cells transduced with control shRNA or shRBPJ. **A**. PCA showing sample clustering by global transcriptional profile. **B**. Heatmap of the top 25 differentially expressed genes between shRBPJ and control samples. Red indicates higher expression and blue indicates lower expression relative to the other condition. **C**. Volcano plot showing differential gene expression between shRBPJ and control samples, with significantly upregulated and downregulated genes highlighted. Significant genes were defined as those with adjusted p < 0.05 and absolute log fold change > 0.5. In the volcano plot, red indicates genes upregulated in shRBPJ and blue indicates genes downregulated in shRBPJ. **D**. GO Biological Process (BP) terms enriched among genes upregulated (top) or downregulated (bottom) in shRBPJ. Negative regulation of cell differentiation is the most enriched pathway (adjusted p < 0.05 and absolute log fold change > 0.5) **E-G**. Heatmaps of gene expression in cells transduced with shRBPJ or control shRNA; red indicates higher expression and blue indicates lower expression relative to the other condition, only genes with an adjusted p < 0.05 are included. **E**. Genes identified as multipotent markers in MCC. Across these gene sets, shRBPJ samples generally show increased expression of differentiation-associated genes and reduced expression of multipotent-associated genes, consistent with a shift toward a more differentiated cell state following RBPJ knockdown. **F**. NE marker genes. **G**. Genes in Termination of RNA polymerase II transcription and related genes.

To further investigate the processes altered by RBPJ knockdown, we performed gene ontology (GO) enrichment analysis as well as gene set enrichment analysis (GSEA) to identify overrepresented biological processes in our significantly DE genes and detect pathways that collectively shift across the ranked gene list. GO Biological Process (BP) terms enriched among upregulated genes in RBPJ silenced cells most notably included negative regulation of cell differentiation and termination of RNA polymerase II transcription (Figure 6D, Supplemental Figure 2A). Other enriched terms were related to wound healing and cell adhesion. In contrast, GO BP terms enriched in genes downregulated in shRBPJ cells were dominated by mitotic and cell cycle-related processes. These results suggest that silencing of RBPJ leads to a loss of the transcriptional activity necessary for cell cycle progression, which is consistent with the reduced proliferation observed in RBPJ silenced WaGa cells. GSEA of genes positively enriched following RBPJ knockdown showed strong enrichment for mitochondrial translation, electron transport chain activity, and ATP synthesis pathways, suggesting that loss of RBPJ is associated with increased mitochondrial and oxidative metabolic programs in WaGa cells (Supplemental Figure 2C). GSEA of the genes negatively enriched following RBPJ knockdown showed strong enrichment for mitotic and cell cycle terms, closely mirroring the trends we saw in the GO analysis of downregulated genes (Supplemental Figure 2D). Together these results indicate that RBPJ knockdown in WaGa cells is associated with broad transcriptional changes marked by reduced mitotic capacity, change in differentiation state, and an altered metabolic state.

### RBPJ knockdown shifts WaGa cells towards a more differentiated transcriptional state

To further assess how RBPJ silencing altered MCC cell state, we examined expression patterns across curated gene sets associated with differentiation, multipotency, neuroendocrine phenotype, Notch signaling, and RNA polymerase II transcriptional regulation (Figure 6E-G and Supplemental Figure 3). First, of the 45 genes upregulated in negative regulation of cell differentiation, 21 of them are PRAME family genes (Supplemental Figure 3A). However, many PRAME family members are poorly characterized individually^64^, making it difficult to functionally interpret changes in these genes. Notably, the best studied family member in cancer, PRAME itself, was not differentially expressed between shControl and shRBPJ cells. Instead, the enrichment of PRAMEF genes, together with the upregulation of DUXA, LEUTX, and ZSCAN4, may be more consistent with activation of a broader ZGA-like process than with negative regulation of cell differentiation. Genes that were differentially expressed in Differentiated versus Multipotent clusters from our initial scRNA-seq data and also extracted as markers of that state from the trained CytoTRACE2 model, were used as markers of the Differentiated or Multipotent state (Figure 6E and Supplemental Figure 3B). Generally, MCC differentiation markers were upregulated in the shRBPJ condition, whereas MCC markers of multipotency were largely downregulated in shRBPJ cells relative to the control. In MCC, MYCL and SOX2 are key stemness factors, and inhibition of SOX2 has been shown to lead to cell cycle arrest and neuron-like differentiation^13,59,65^. Here, we observe a downregulation of SOX2 in shRBPJ cells relative to the control, along with an upregulation of MYC and MYCN, indicative of a decrease in MYCL^22^. Our previous work has also shown that WNT5A expression, which here is overexpressed in shRBPJ cells, is negatively correlated with MCC markers^42^. Together, these data support the conclusion that loss of RBPJ shifts WaGa cells away from a more stem-like program and toward a more differentiated transcriptional state.

Additional gene sets provide context surrounding this transition. Some Notch related genes were perturbed as expected in shRBPJ WaGa cells e.g. NOTCH3, NOTCH4, and DLL1^66^. However, some of the transcriptional changes more closely align with expression patterns observed in glioblastomas (Supplemental Figure 3C)^32^. For example, in glioblastoma, RBPJ was found to be a direct regulator of FOXM1, CCNA2, and KRAS, which control brain tumor-initiating cell proliferation, and when RBPJ was knocked down, expression of these genes decreased, which is consistent with our observations. Additionally, in glioblastoma, silencing of RBPJ had little effect on canonical Notch targets including HES1^32^, which we see upregulated in shRBPJ cells. In other contexts, such as autoimmune eye disease, RBPJ knockdown leads to a significant decrease in HES1^67^. Neuroendocrine related genes also showed marked changes following the knockdown of RBPJ (Figure 6F). The well-known MCC markers POU4F3^42^ and INSM1^68^ were downregulated in shRBPJ cells, whereas ASCL1, which is a marker of lung neuroendocrine markers but is not normally expressed in MCC^69,70^, was upregulated, indicated a change in cell differentiation and lineage specificity. Finally, we observed changes in the expression of genes related to RNA polymerase II transcription and the cell cycle consistent with what has been previously observed in glioblastoma (Figure 6G). The SSU72-like family, known to be involved in termination of RNA polymerase II transcription^71^, is upregulated with RBPJ silencing, whereas POLR2A, the main subunit of RNA Pol II, and CDK9, kinase subunit that promotes transcriptional elongation^72^, are both downregulated. We also observe an upregulation of CDKN1A, the cyclin-dependent kinase inhibitor p21, and downregulation of cyclin B1 in the RBPJ knockdown cells, highlighting a slowing of cell cycle progression^73^. The downregulation of BIRC5, the oncoprotein and apoptosis inhibitor Survivin^74^, in shRBPJ cells is consistent with our observation that these cells exhibit reduced growth in a manner that could be clinically useful. We also see reduced expression of AURKA and AURKB in shRBPJ cells. Inhibition of AURKB has been found to induce apoptosis in MCC cells and inhibit tumor growth in xenograft models^75^, possibly explaining in part the decrease in cell growth rate we observe when silencing RBPJ. Taken together, our data demonstrate not only a shift of MCC cells away from a more stem-like state, but also a loss of MCC markers, and a reduction in proliferation following RBPJ knockdown, reminiscent of the sensitivity to RBPJ inhibition reported in glioblastomas.

## Discussion

In this study, we use probabilistic Boolean networks to model lineage plasticity in MCC and predict key regulators of differentiation, including RBPJ, that when silenced, reprogram the cell to a more differentiated and treatable state. Working across two independent datasets, we identified four potency states in MCC VP tumor cells: Differentiated, Unipotent, Oligopotent, and Multipotent. We then implemented a modified version of the BooleaBayes algorithm to model gene-level perturbations of our network, revealing RBPJ as the most significant regulator of the MCC differentiation network. In order to capture more system-specific regulatory relationships, we also established a workflow to expand the regulatory network using ATAC-seq inferred target genes. This led us to generate two additional networks, both of which also yielded RBPJ as the most significant regulator of differentiation, demonstrating the robustness of these simulations. By silencing RBPJ in MCC cells, we discovered that this leads to dysregulation of differentiation pathways and a substantial MCC-specific decrease in growth rate. Beyond a decrease in both the NE and multipotency signature, knockdown of RBPJ in MCC also caused transcriptional changes in Notch pathway targets and cell cycle regulation concordant with previous studies on glioblastoma. Our work provides a reproducible workflow for modeling cell-state transitions *in silico* to prioritize TFs that regulate differentiation, and it also contributes to our understanding of RBPJ in MCC, highlighting it as a potential novel target for treatment in this disease.

The Notch pathway is a juxtracrine signaling mechanism, in part responsible for cell fate determination through lateral inhibition^31,76^. For such functions, there is a complex coordination of activation receptors by different ligands, leading to activation of different transcriptional targets under different conditions. RBPJ, often thought of as a mediator of Notch signaling, has different roles itself, acting as an activator or inhibitor under separate circumstances^32,34,77^. As may be expected, cancer further complicates this model as Notch is implicated to be a tumor suppressor in certain tumors and an oncogenic factor in others. While inhibition of RBPJ leads to suppression of tumor growth in glioblastomas, in primary human dermal fibroblasts it activates a cancer-associated fibroblast phenotype^60,61^. This is congruent with our increased growth rate of HDFn cells and decreased growth rate of WaGa cells. Thus, we hypothesize that RBPJ has a similar role as in glioblastomas, which implies that it is involved in the maintenance of MCC cells through Notch-independent mechanisms. Consistent with this hypothesis, our data show that HES1 is upregulated in the absence of RBPJ, and the expression of RBPJ is correlated with FOXM1, OLIG2, SOX2, and CHD7. It has been reported that FOXM1 and OLIG2 are activated by RBPJ independent of Notch signaling, while SOX2 and CHD7 interact with RBPJ in cancer-specific pathways^32,78^. Future work is necessary to characterize these interactions in MCC and determine if the tumors are adopting similar neural stem cell pathways.

In glioblastomas, MYC was found to directly regulate expression of RBPJ and CDK9 was discovered to bind to RBPJ to regulate transcription. Cyclin-dependent kinase 9 (CDK9) is a core subunit of positive transcription elongation factor b (P-TEFb), which is responsible for the regulation of RNA polymerase II elongation^72^. As such, many cancers are associated with an increase in function of CDK9 through mechanisms such as P-TEFb-dependent amplification of gene expression by MYC^79^. In glioblastoma, it has been proposed that MYC activates expression of RBPJ, and RBPJ then recruits CDK9 to activate expression of its target genes^32^. Our data suggests a similar interplay may be present; in our RBPJ silenced cells we observed a downregulation of CDK9 and POLR2A with an upregulation of RNA pol II transcription termination factors, leading us to believe there is a decrease in transcriptional activity. This along with the dysregulation of other cell cycle factors leads to the decreased growth rate of WaGa cells we observed. Notably, however, MYC expression is not typically amplified in MCC^13^. Instead, L-Myc is a driving factor of tumorigenesis and dedifferentiation in MCC^59^, and the expression of c-Myc is downregulated, with L-Myc and c-Myc appearing to play opposing roles. Consistent with this, we found that RBPJ knockdown leads to upregulation of MYC and downregulation of MYCL. Taken together, it is possible that L-Myc, not c-Myc, drives RBPJ expression in MCC. Future studies are needed to elucidate the regulatory targets of RBPJ in MCC, evaluate binding to CDK9 and its role as a transcriptional regulator, and identify the upstream factors that drive RBPJ activity.

Similar to any dynamic gene regulatory network model, our workflow can only partially capture MCC-specific regulatory data. The BooleaBayes package was developed for small, strongly connected networks where every node has at least one target and source. As an example, the literature-based prior network from Wooten et al.^28^ had 16 nodes with an average of 6.4 neighbors per node, while our prior network of 61 nodes, consisting mostly of edges inferred by DoRothEA, only had an average of 2.5 neighbors per node. This meant that the computational overhead increased exponentially for an attractor search, and those found were less biologically relevant as the presence of many nodes with no targets meant that there were more ways the system could end up stable as they do not ‘feed back’ into the network. To try to better model the complexity of the network, we increased the sensitivity of our network prior by adding MCC specific edges derived from ATAC-seq data. This increased the average number of neighbors to around 14 neighbors per node. Interestingly, the regulators predicted to drive cell differentiation were consistent between three different versions of the prior network, suggesting that our predictions are driven by a core regulatory circuit involving known regulatory interactions. These models appear to be sufficient for capturing important changes in cell differentiation through single perturbations of transcription factors. In contrast, when we applied SCENIC^80^ to model the regulatory network, it was unable to assign RBPJ to a regulon and therefore could not identify it as a regulator of multipotency. Instead, the TFs with biggest differences in regulon activity score between the differentiated and multipotent clusters included factors such as KLF8, RFX1, and DRGX, identifying more general differences of these clusters in the cell cycle rather than the underlying mechanisms of this transition (Supplemental Table 1). Thus, the BooleaBayes model was able to capture regulatory mechanisms beyond SCENIC.

Another limitation of our study is that currently the dynamic simulations are synchronously updated, meaning all TFs update their state at the same time. To better represent a real biological system, we could incorporate an asynchronous updating mechanism, which may yield more realistic results^58^. For improved interpretability of the simulations, future work should also focus on refining the attractor state search for large networks. While the current algorithm is exponentially more time and memory intensive with increasing network size, we can optimize the algorithm to minimize use of both by selectively precomputing and mapping binary states, thereby reducing the number of mappings and duplicate searches. Additionally, we can improve efficiency by searching for attractor states around pre-defined points in state space for downstream simulations as opposed to searching through the entire landscape of possibilities.

Altogether, our work demonstrates that combining probabilistic network modeling and high-throughput genomic data can uncover key mechanisms of cell state regulation in Merkel cell carcinoma. Inhibiting the master regulators driving multipotent cells in MCC tumors could yield new treatment strategies that target the most aggressive cells and lead to clinical benefit.

## Acknowledgements

This work was supported by NIH grant R01CA251729 to MP.

## Data availability

The data generated in this study will be made available in Gene Expression Omnibus upon publication.

---

**Supplemental Figure 1.**
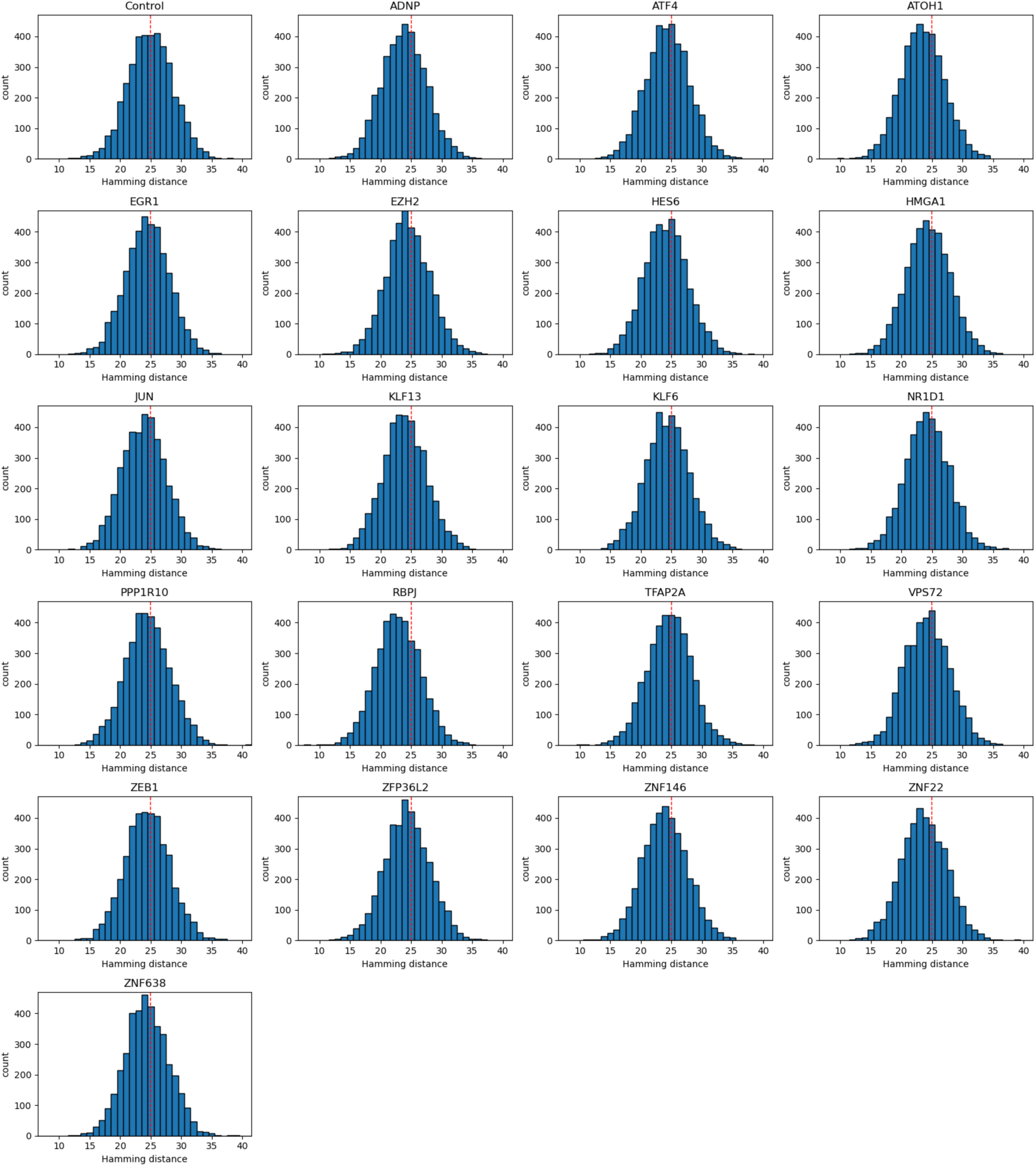
Histograms showing, for each significant gene, the distribution of cell counts at each Hamming distance from the reference state for the final 2 time steps. Red line is the average final distance for the control.

**Supplemental Figure 2.**
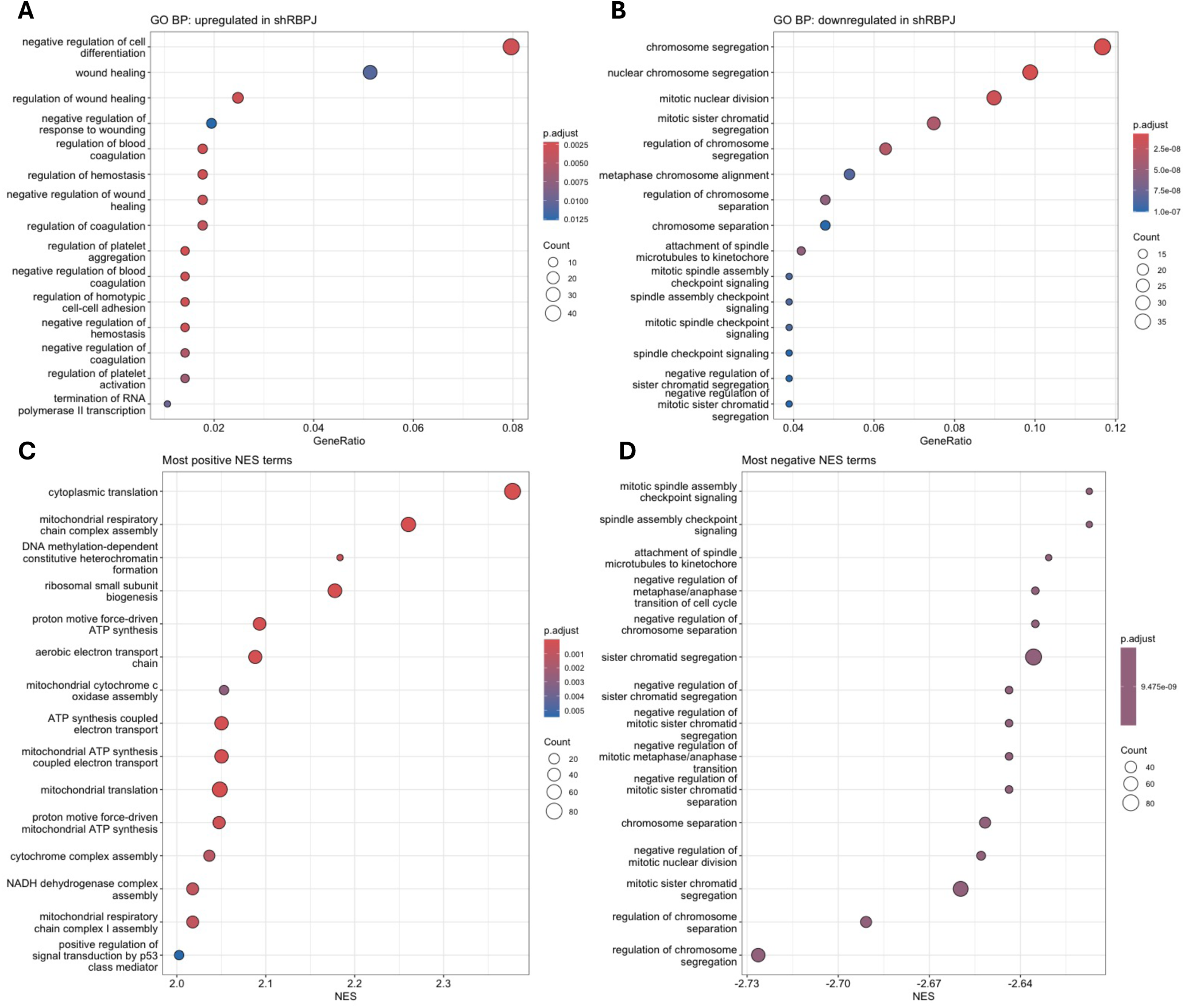
Gene ontology (GO) and gene set enrichment analysis (GSEA) were performed on bulk RNA-seq data from WaGA cells transduced with control shRNA or shRBPJ. **A, B**. GO Biological Process (BP) terms enriched among genes upregulated (A) or downregulated (B) in shRBPJ. Negative regulation of cell differentiation is the most enriched pathway (adjusted p < 0.05 and absolute log fold change > 0.5). **C,D**. GSEA of positively (C) and negatively (D) enriched terms in shRBPJ relative to the control. Dot size represents the number of genes contributing to each term, and color indicates adjusted p value.

**Supplemental Figure 3.**
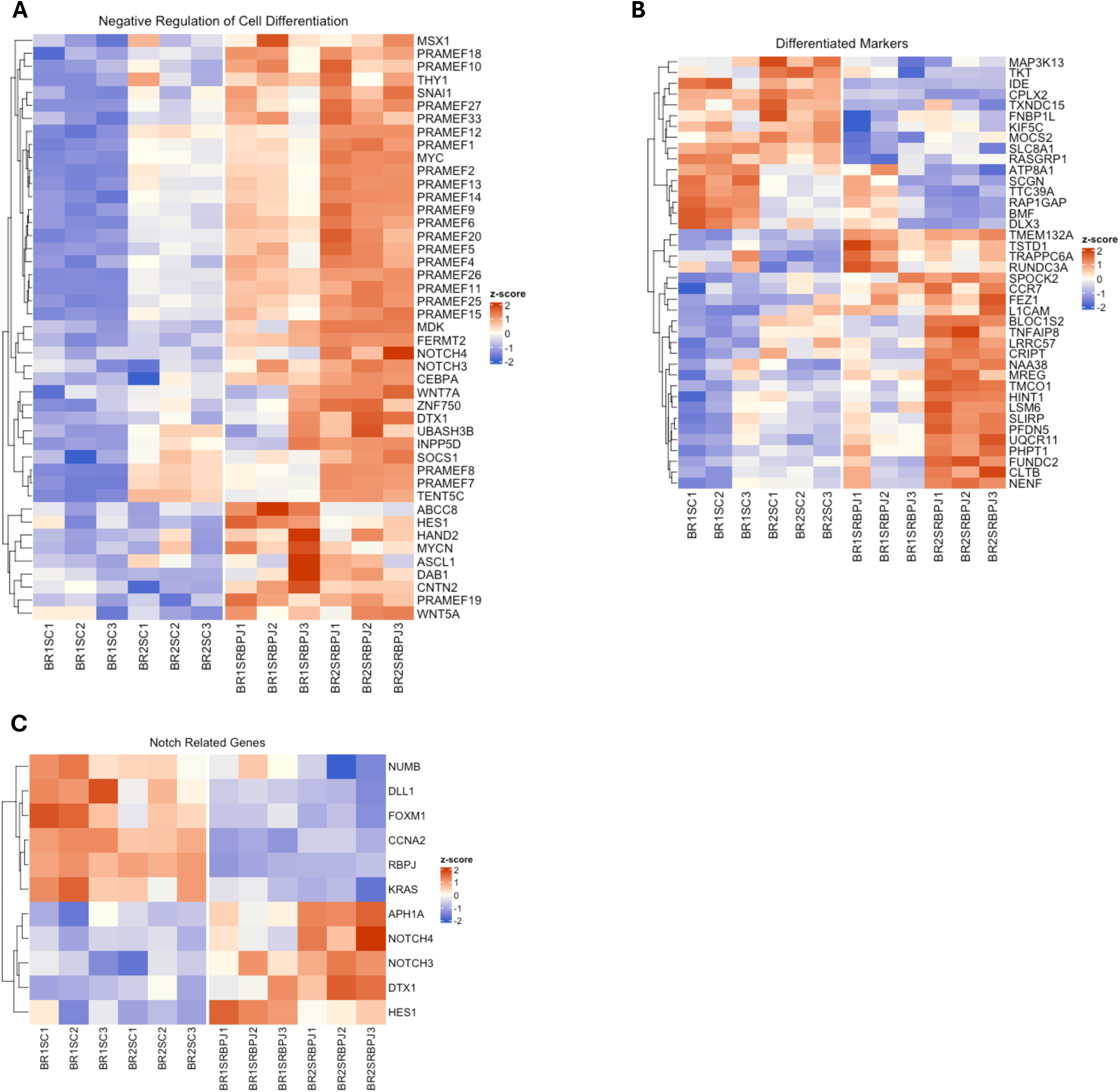
Red indicates higher expression and blue indicates lower expression relative to the other condition, only genes with an adjusted p < 0.05 are included. **A**. Heatmap of genes in the Negative Regulation of Cell Differentiation set. **B**. Genes identified as markers of differentiation in MCC. **C**. Notch related genes and targets of RBPJ.

**Supplemental Table 1.** Top enriched transcription factors using SCENIC. TFs were sorted by the absolute difference in mean AUC between the cells in the differentiated and multipotent clusters.

|  | TF | maxNES | n motifs | n targets | meanAUC | AUC diff | AUC multi | deltaAUC | abs_deltaAUC |
| --- | --- | --- | --- | --- | --- | --- | --- | --- | --- |
| 1 | KLF8 | 3.23015419 | 1 | 4 | 0.02805066 | 0.05844916 | 0.00592221 | 0.05252694 | 0.05252694 |
| 2 | RFX1 | 3.41852355 | 2 | 12 | 0.06330444 | 0.02405523 | 0.06035241 | -0.0362972 | 0.03629718 |
| 3 | DRGX | 3.60026687 | 1 | 12 | 0.070119 | 0.09096005 | 0.06099841 | 0.02996164 | 0.02996164 |
| 4 | PBX1 | 3.48908726 | 1 | 7 | 0.08226531 | 0.06244833 | 0.09035349 | -0.0279052 | 0.02790516 |
| 5 | RFXANK | 4.28674673 | 4 | 19 | 0.02946144 | 0.04631392 | 0.01994633 | 0.02636759 | 0.02636759 |
| 6 | POU4F3 | 3.17971454 | 1 | 7 | 0.10959221 | 0.09868368 | 0.12229332 | -0.0236096 | 0.02360965 |
| 7 | CREB3 | 3.57624037 | 2 | 20 | 0.02423136 | 0.04008886 | 0.01654524 | 0.02354362 | 0.02354362 |
| 8 | ZBTB11 | 3.85669929 | 3 | 259 | 0.04918051 | 0.02829331 | 0.05005175 | -0.0217584 | 0.02175844 |
| 9 | LEF1 | 3.72416023 | 2 | 13 | 0.01059016 | 0.02272143 | 0.00119545 | 0.02152598 | 0.02152598 |
| 10 | RUNX3 | 4.7252653 | 5 | 212 | 0.02855995 | 0.04144959 | 0.02015524 | 0.02129436 | 0.02129436 |
| 11 | FOXA3 | 4.29853435 | 5 | 22 | 0.02848343 | 0.04341365 | 0.02223105 | 0.0211826 | 0.0211826 |
| 12 | CEBPG | 9.72620609 | 9 | 454 | 0.07895646 | 0.06440125 | 0.08531569 | -0.0209144 | 0.02091445 |
| 13 | EOMES | 3.66568038 | 3 | 43 | 0.01123588 | 0.02307117 | 0.00217437 | 0.0208968 | 0.0208968 |
| 14 | GATA3 | 3.85440814 | 7 | 200 | 0.01677948 | 0.02826693 | 0.00827143 | 0.0199955 | 0.0199955 |
| 15 | IKZF2 | 4.56983899 | 14 | 318 | 0.02228486 | 0.03397424 | 0.01462555 | 0.0193487 | 0.0193487 |
| 16 | SPIB | 5.6747898 | 10 | 311 | 0.03068172 | 0.04215645 | 0.02281556 | 0.01934089 | 0.01934089 |
| 17 | STAT6 | 3.90639054 | 12 | 109 | 0.01106567 | 0.02139688 | 0.00302768 | 0.0183692 | 0.0183692 |
| 18 | MAFB | 3.37088888 | 2 | 12 | 0.0866556 | 0.07683675 | 0.09443064 | -0.0175939 | 0.01759389 |
| 19 | ZBTB21 | 3.1737365 | 2 | 325 | 0.04713317 | 0.03233342 | 0.04872168 | -0.0163883 | 0.01638826 |
| 20 | NFATC2 | 5.65986408 | 4 | 155 | 0.01240603 | 0.02134916 | 0.00574012 | 0.01560905 | 0.01560905 |

